# Octopamine and Tyramine Modulate Fictive Motor Program Competition in *Drosophila* larvae

**DOI:** 10.1101/2025.08.17.670699

**Authors:** William V. Smith, Stefan R. Pulver

## Abstract

Dynamic interactions amongst competing motor programs shape behavioural output. Here, we use bath application of biogenic amines octopamine (OA) and tyramine (TA) to explore operating principles of competition amongst central pattern generating (CPG) networks controlling locomotion in 3 ^rd^ instar *Drosophila* larvae. OA and TA share the same synthesis pathway and are contained in overlapping sets of neurons. Bath application of OA promoted fictive forwards locomotion, suppressed backwards locomotion, and induced bouts of fictive head sweeps during wash periods that were proportional to the promotion of forward waves. In contrast, TA, the conceptualised inhibitory antagonist to OA, promoted collisions and overlap of motor programmes, activity-dependent inhibition, and bouts of silence during wash periods. Co-application of OA and TA revealed bimodal state-dependent modulation with forward bias promotion exclusively if the pre-application network was not already forward dominant. Overall, our work provides insights into how adrenergic-like systems can modulate dynamics of motor competition among interacting CPGs within locomotor networks.

## Introduction

Effective locomotion depends on neural circuits that achieve a dynamic balance between stability and flexibility. Motor behavior arises not only through the initiation of specific motor programs but also through the suppression of pattern generation circuits that would induce alternative motor programs – motor competition [1], [2], [3], [4], [5], [6], [7]. Coordinated locomotion requires the sustained activation of central pattern generators (CPGs) in concert with precisely timed inhibition of premotor networks that drive opposing CPG activity. Disruption of inhibitory interactions between CPG networks can therefore lead to uncoordinated or dysrhythmic motor output [1]. To preserve both stability and adaptability, motor systems must support the initiation and maintenance of locomotor activity while retaining the capacity to transition between alternative motor programs as conditions change. Conceptually, distinct pattern-generating networks can be viewed as competing for control within a temporally and spatially constrained neural landscape to produce context-appropriate motor output. Because motor adaptability is essential for survival—for example, during rapid escape responses—understanding how motor circuits modulate and arbitrate such motor competition may reveal general principles of neural flexibility and resilience. Although the circuit motifs underlying motor program selection and coordination are increasingly well characterized (e.g., [7], [8], [9], [10], [11], [12]) the neuromodulatory mechanisms governing competitive selection among motor programs remain comparatively unresolved.

The adrenergic-like system in *Drosophila* provides a powerful testbed to evaluate motor competition by enabling controlled, acute induction of motor bias. The *Drosophila* adrenergic-like system is composed of octopamine (OA) - an invertebrate analog to adrenaline - and tyramine (TA) [13], [14]. The biosynthetic pathways of OA/TA, their receptor pharmacology [15], [16] and the expression patterns of OA and TA neurons are well studied in multiple invertebrate species [15], [17], [18], [19], [20], [21], [22], [23], [24], [25], [26], [27], [28], [29]. The adrenergic-like system has been implicated in a variety of fundamental behaviours including learning and memory [15], [30], social behaviours like aggression [31], [32], [33], motor initiation and modulation [14], [21], [34], [35], [36], including fundamental processes underlying action like energy metabolism [37], [38], synaptic plasticity [39], [40], and activity-mediated adaptation [41], [42]. The adrenergic-like system is consequential to invertebrate movement especially in its capacity to modulate selection of motor programs based both internal states and external sensory drives.

Traditionally, OA and TA are considered neuromodulators with opposing actions and outcomes [43], [44], [45]. Namely, OA has a well-documented promotive and facilitatory role in locomotion [21], [34], [35], [35], [35], [36], [46], [47], [48] while TA appears to have an overall inhibitory effect on locomotion [18], [28], [43], [45], [49] across invertebrates. However, the functional relationship between OA and TA’s action is not always exclusively antagonistic nor definitely excitatory/inhibitory. For instance, the mutual-antagonistic relationship of OA/TA appears to be conditional on their mutual relative concentrations [44]. Further, OA and TA are not always generally excitatory or inhibitory, respectively. For example, octopamine can have inhibitory effects on CPG activity through inhibition of CPG command interneurons [50], can inhibit electrically stimulated locomotion [51], and can inhibit synaptic transmission at the *Drosophila* larval neuromuscular junction [52]. Indeed, different OA receptors are linked to either stimulatory (Octβ2R) or inhibitory (Octβ1R) cAMP/CREB pathways, which form the basis of self-regulated octopaminergic synaptogenesis at the larval muscular junction [39], [40]. On the other hand, TA can have excitatory effects on neural networks underpinning behaviour. For example, TA can evoke depolarisation through an indirect DA-mediated pathway in rat subthalamic neurons [53], at low concentrations in hyperpolarized states increase the amplitude of locust oviduct excitatory junction potentials stimulating rhythmic contractions [54], and facilitate sensitisation by delaying cocaine-induced responsive hypoactivity [55]. Nonetheless, a significant body of work points to a binary perspective on OA/TA within locomotory systems. The binary appreciation of neuromodulation may be a result of experimental approaches that may create developmental compensatory effects (e.g., RNAi, genetic ablation) [56]. By evaluating adrenergic-like signalling through an acute experimental approach we may gain insights into moment-to-moment modulatory effects that are more ecologically valid [57], [58], [59].

The role of the adrenergic-like system in modulating action selection and motor pattern competition has been extensively studied in *C. elegans* through tyraminergic RIM neurons [60], [61], [62] which has instantiated a fundamental adrenergic-like motif (i.e., “master-and-slave”) in *C. elegans* locomotion. However, while the role of OA/TA has been studied in *Drosophila* with reference to specific motor program initiation and coordination [63], limited work has been conducted on their coordinated role to generalisable motor competition. Through studying the effects of acute modulation on action selection and motor competition, there is an opportunity to gain a more robust, nuanced, and applicable understanding of OA/TA neuromodulatory antagonism.

Here, we study OA and TA modulation of locomotor activity in *Drosophila* larvae to better understand the neural architecture behind CPG coordination and competition. We demonstrate how bath-applied exogenous OA and TA induce acute changes in CPG activity during, and after bath application. Specifically, OA induces elevated fictive forward locomotion in isolated central pattern generators. In contrast, exogenous TA induces increased overlapping fictive patterns with activity-dependent inhibitory effects on general fictive activity. Both neuromodulators also induce long lasting effects following bath application that reflect activity changes in the network. Co-application of OA and TA reveal state-dependent promotion of bias antagonistic to the fictive basis in the pre-application period. Overall, we demonstrate how reactive motor competition could be an informative way to conceptualise motor diversity and neuromodulation and how the insect adrenergic-like system is a valuable tool for understanding the secondary effects of modulating motor competition.

## Methods

### Animal Rearing & Dissection

*Drosophila* were raised at 18-25◦C in incubators on an approximate 12-hour light-dark cycle. All experiments were conducted with feeding 3 ^rd^ instar *Drosophila* larva (2-3 days post hatching). The central nervous system (CNS) of *Drosophila* was isolated as reported in [64], [65]. Activity of isolated CNSs was recorded with Baines External Saline (BES) submersion containing (in mM) 135 NaCl, 5 KCl, 2 CaCl_2_, 4 MgCl_2_, 5 TES, and 36 sucrose, pH 7.15 [66]. All CNS preparations were excised from *Drosophila* larval bodies using a dorsal incision across the larval body using fine scissors with body walls pinned into a Sylgard-lined Petri dishes using fine tungsten wires (California Fine Wire, Grover Beach, CA). All projecting nerves from CNSs were cut, CNSs were then removed, and posterior nerves and peripheral tissues attached were pinned using fine tungsten wires (see [1], [64]).

### Ca^2+^ Imaging with Tyramine & Octopamine Perfusion

The effect of exogenous octopamine bath application on calcium signals in motor neurons was assessed in isolated CNS feeding 3^rd^ instar *Drosophila* larvae using a genetic line in which GCaMP6m is driven in glutamatergic neurons by the OK371-GAL4 line. The posterior nerves of isolated CNS preparations were pinned to Sylgard-lined Petri dishes using thin tungsten wires. Saline and OA and/or TA were applied during imaging using a gravity-based perfusion system with a consistent drip inflow across preparations and removed continuously using a KCP3-B10W peristaltic pump (Kamoer 60-120 RPM). Across all preparations, saline was bath applied onto the preparation for at least 10 minutes followed by 10-minute OA or TA application then < 20 minutes saline reapplication to wash away octopamine/tyramine from the Sylgard-lined Petri dish. All OA concentration ranges (5 - 400μM) were generated from a diluted 0.02M master mix (O0250-1G Sigma Aldrich). All TA concentration ranges (5 - 400μM) were generated from a diluted 0.01M master mix (T90344-5G Sigma Aldrich). Images were collected using WinFluor (University of Strathclyde, UK) at 10fps and post-processing using ImageJ [67]. All isolated preparations were recorded using a UPlanFLN Olympus 10x, 0.3nA.

### Electrophysiology with Tyramine & Octopamine Perfusion

The effect of OA and TA perfusion on fictive motor activity was studied using extracellular electrophysiology on segmental nerve roots in isolated CNS from feeding 3^rd^ instar *Drosophila* larvae. Across all conditions, the activity of either thoracic nerve root (T3) and/or posterior abdominal nerve root (A6-A8) was measured by suction electrode recording (AM System Differential AC Amplifier Model (#59721, 1700), 1-100kHz cut-off frequency allowance). Borosilicate glass capillaries were pulled using an electrode puller (Narishege, Japan), and tapered tips were broken to produce suction electrodes that fit tightly around single nerve roots. All neuromodulators were bath applied.

All electrophysiology recordings and light stimulations were made using LabChart 8.0 software (AD Instruments, Colorado Springs, CO) and post-processed in Spike2 (Cambridge Electronic Design, Cambridge, UK) and Python3. Post-processing involved rectification and smoothing using a consistent 0.2s time interval. All rectified and smoothed traces were imported into DataView where event peaks and 20% on-off duration were determined by Hill’s Valley analysis. For analysis of nerve bursting effects of TA bath application, the frequency of activity in A6 was determined within bursting bouts. Bursting bouts were defined as blocs of activity flanked by at least 5 seconds of no peak activity. Visualisation and further analysis were conducted in Python3 (see repository code).

### Classifying Fictive Activity

Fictive activity was classified based on detecting peak in motor neuron calcium signals and using strict definitions of intersegment peak activity. The motor dynamics within the isolated VNC preparation were classified using Hill’s Valley thresholding to determine peaks within motor activity in each abdominal and thoracic ganglion segment alongside established definitions for each fictive behaviour [1], [64], [65] (Figure 1). Succinctly, fictive forward waves and fictive backward waves were determined by the existence of sequential Hill’s Valley-determined peaks proceeding from either posterior-to-anterior or anterior-to-posterior in all segments, respectively. Fictive headsweeps were determined by computing a difference trace (left – right) in the T3 segment and determining anterior asymmetric activities as peaks or troughs that exceeded ±1% absolute threshold difference. Turning behaviours were defined as an anterior asymmetric activity immediately followed by a fictive backward event [68]. Finally, anterior symmetric (i.e., anterior burst) or posterior symmetric (i.e., posterior burst) were determined by peak detection of the left and right trace in T3-A2 and A6-8 respectively. Some anterior burst and posterior burst fictive activities are in varying numbers of anterior and posterior segments thus any peak detection in both the left and right trace within 5s of each other was determined as symmetric activities of each definition. Fluorescence traces were converted to ΔF/F by estimating a dynamic baseline for each ROI using a 20th-percentile filter over a 500-sample moving window. ΔF/F was calculated as (F − F_0_) / F_0_. Smoothed traces were generated from the ΔF/F signal using a second-order Savitzky–Golay filter with a 23-sample window. In the calcium dataset, Hill’s Valley peak detection was performed after baseline correct (df/f) and smoothing within DataView [1]. In the electrophysiology dataset, raw electrophysiological traces were smoothed in Spike2 thereafter DataView was used to determine peaks in the anterior and posterior trace akin to the calcium dataset analysis pipeline. Specifically, the determination of fictive forward vs backward activity was determined by the minimum peak-to-peak time between the initiating-vs-terminating segment. Specifically, if the peak time difference between T3-to-A8 was shorter than the peak time between A8-to-the-next-T3 peak then the activity peaks were classified as fictive backward events rather than fictive forward events. If the activity peak differences exceeded 3s, the activity peaks were either reported as anterior bursting or posterior bursting events in T3 or A8, respectively. In cases of ambiguity, the fictive events were not recorded. Manual corrections – deletions or additions of peaks – were performed, when necessary, in line with the strict fictive definitions highlighted above.

**Figure 1.**
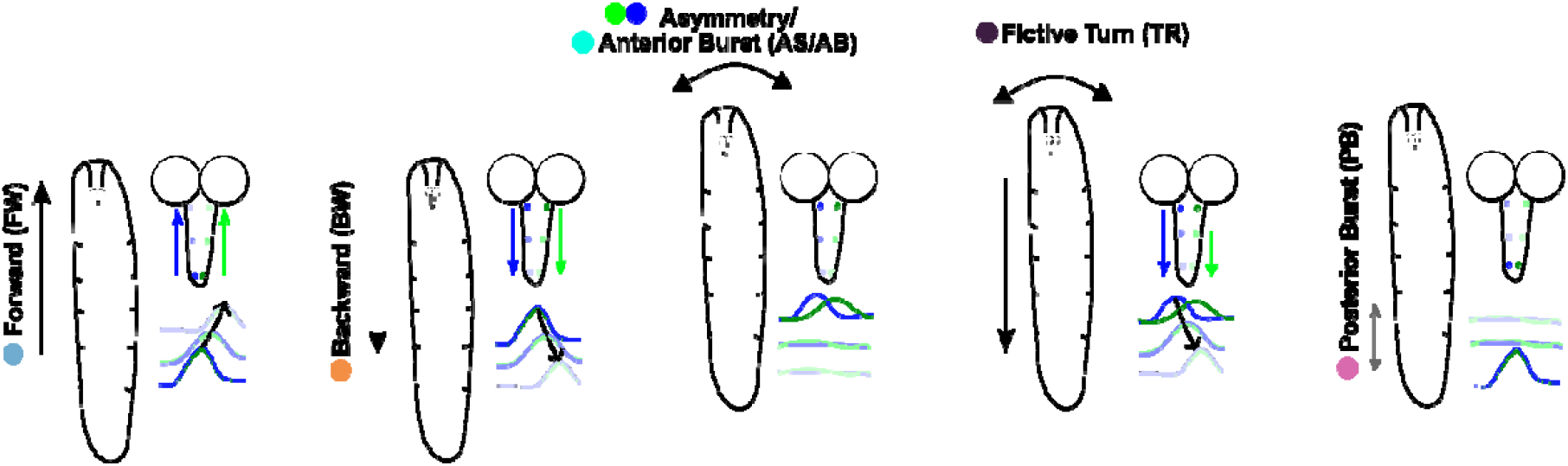
Fictive Behavioural Categories. Fictive behaviour of the *Drosophila* larva is classified by different motor neuron activity sequences within the isolated ventral nerve cord (VNC) (as originally noted in [64]). Fictive activity patterns are shown alongside analogous intact behavioural patterns. Forward crawling consists of segment contractions in a posterior-to-anterior direction induced by a posterior-to-anterior sequence of motor neuron activity. Backwards crawling consists of segment contractions in an anterior-to-posterior direction induced by an anterior-to-posterior sequence of motor neuron activity. Headsweep behaviour – angular sweeping of the anterior regions of the intact animal – is induced by asymmetric activation of either the left or right side of motor neurons in the thoracic ganglion. Head crunches/raises are likely induced by anterior symmetric activation of motor neurons in the thoracic ganglion. Turning behaviours consists of a headsweep which remains unresolved while the larva retreats induced by initial asymmetric activity in motor neurons in thoracic ganglion with symmetric anterior-to-posterior progression of motor neurons in the abdominal ganglion. Posterior crunches consist of posterior segment contractions induced by symmetric activity in the motor neurons in the posterior segments of the VNC. All subsequent calcium imaging analysis categorises sequential motor activity in either of the six fictive patterns: fictive forward (FW), fictive backward (BW),anterior asymmetric (AS), turns (TR), anterior bursting (AB), or posterior bursting (PB) events.

Activity, silence, and overlap incidence for perfusion analysis was conducted in DataView with strict definitions. The duration of total fictive activity was determined through the same Hill’s Valley peak detection algorithm used to initially categorise fictive activity patterns. Specifically, the total duration of fictive activity within a condition time was calculated as the 80% margins before and after peaks in calcium activity from all segments that exceeded 1% standard deviation above the baseline signal. Silence was as the percentage of time when no calcium trace showed activity ≥ .01 df/f in any part of the VNC compared to the total recording period. Fictive overlap was defined as the existence of the start, or progression, of a fictive behavioural program in the network before the definitional completion of a prior fictive program. For instance, a fictive forward wave would be overlapping a fictive backward wave if the A8 initiation peak of the fictive forward wave occurring before the T3 completion peak of the fictive backward wave. Exclusively fictive wave motor programs were considered in the definition of overlapping fictive events.

### Analysis, Visualisation, & Statistical Evaluation

All calcium signals were initially analysed in DataView (version 12.2.2) [69], [70] using Hill’s Valley Analysis for peak detection. All data were visualised in Python3 using custom scripts available in the repository supplied by the authors. All statistical tests were performed using non-parametric packages within Python (see repository scripts and report tests in the results sections).

#### Analysis of Bimodality

Bimodality in the simultaneous perfusion dataset was determined by KMeans clustering of % silence during simultaneous co-application of octopamine and tyramine, subject to between-clusters validation through a Mann Whitney U test, whose robustness was validated by a 2-component Gaussian Mixture Model.

The structure of calcium activity traces of all motor neuron pools during co-application of OA and TA that resulted in low-amplitude, no detectable fictive rhythms was evaluated through 4 metrics. Firstly, the concentration of power spectra density (PSD) in a specific frequency band relative to the total power (**1**). Secondly, peak prominence which quantifies how much dominant spectral peak stands out from surrounding frequencies capturing the sharpness of dominant rhythms (**2**). Thirdly, signal-to-noise ratio-like metric (SNR-like) which compares the power in the dominant and power outside the band (**3**). Fourthly, spectral entropy which measures how evenly-distributed power is across frequencies (**4**).

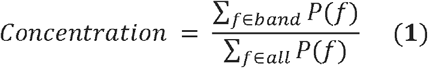

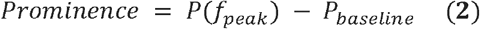

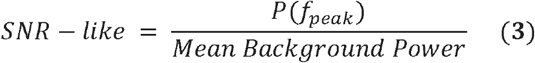

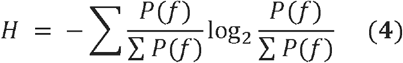

Exploration of pre-application state-dependent effects upon octopamine and tyramine co-application was evaluated through systematic evaluation of fictive-related metrics (see Supplementary Table 2). The following metrics were evaluated: i) proportion (%) of fictive events, ii) pre-application relative silence (see Classifying Fictive Activity), iii) the probability that behavioural state at once instance is the same at the next instance (i.e., stickiness), iv) the number of transitions between one fictive event and another within the recording time (i.e., switch rate). For each computed metric, univariate statistical tests (Mann-Whittney U and Cliff’s delta) were applied to assess metric’s distributional separation. The predictive metric was determined by each metric’s ROC-AUC with AUC results strongest for forward-backward-related metrics (i.e., FW:BW). Lasso regression was performed to determine robust directional discriminators. Given the strongest predictors for promotive and suppressive regimes were all forward-backward metrics and highly intercorrelated, we applied principal component analysis (PCA) to compress directional discriminators to a dominant axis denoted as the directional commitment axis (DC1) for OA and TA co-application which is expressible as (**5**) where *z*(·) represents standardised values, *ω* _*i*_ are PCA loadings, and signs are chosen so higher DCI is reflective of more forward bias. Values are found in Table 1.

**Table 1.** Values from PCA for DCI.

| $\omega_i$ | Value |
| --- | --- |
| 1 $\{z(FW\%)\}$ | 0.381 |
| 2 $\{z(BW\%)\}$ | 0.379 |
| 3 $\{z(FW\% - BW\%)\}$ | 0.412 |
| 4 $\{z\left(\frac{FW}{BW}\right)\}$ | 0.294 |

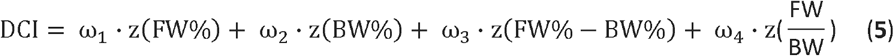

### Sustainability Statement

Through using WillCO_2_st [71], [72], we determined that all experimental work presented here generated 798.23g CO_2_e from the South Scotland UK National Grid.

## Results

### Bath Application of OA induces Fictive Forward & Long-lasting Anterior Asymmetric Activity

Exogenous application of OA to intact invertebrates has a well-reported role as a promoter of locomotion aligned with fight-or-flight responses [15], [21], [73]. Here, we explored how biased induced by exogenous application of OA shapes the diversity of fictive rhythms.

Exogenous application of OA shifted the isolated *Drosophila* ventral nerve cord (VNC) from producing a diverse range of fictive behaviours to predominantly fictive forward and anterior asymmetric motor patterns (Figure 2A). Application of OA concentrations ≥100μM significantly promoted fictive forward activity (100 μM 5.4±2.0 m^-1^ p = .003; 200 μM 6.5±3.2 m^-1^ p < .001; 400 μM 14.9±6.4 m^-1^ p = .004) compared to saline pre-application (100 μM 0.7±0.4 m^-1^; 200 μM 0.7±0.7 m^-1^; 400 μM 0.8±0.7 m^-1^) (Figure 2Bi). At OA concentrations ≥200μM, the induction of fictive forward bias came at the expense of fictive backward motor activity which was significantly reduced (200 μM 0.2±0.6 m^-1^ p = .005; 400 μM 0.6±0.7 m^-1^ p = .004) compared to saline pre-application (200 μM 1.4±0.5 m^-1^; 400 μM 2.3±0.9 m^-1^) (Figure 2Bii). Anterior asymmetric events were significantly promoted at higher OA concentrations 400μM (Figure 2Bv). Importantly, anterior asymmetric promotion was not during the 10-minute OA application but only upon OA wash following the OA-induced fictive forward bias (Figure 2A, Bv). The promotion of anterior asymmetric events by OA peaked 9.2 minutes post the start of the OA wash period (Figure 2E). By considering the change in the probability of transition between different fictive motor programs, we can see the OA-induced collapse in diversity of fictive motor programs is due to universal bias away from all other non-forward motor programs (Supplementary Figure 1B). Overall, exogenous application of OA biases the isolated network towards a forward mode with a post-application bias to anterior asymmetry activity.

**Figure 2.**
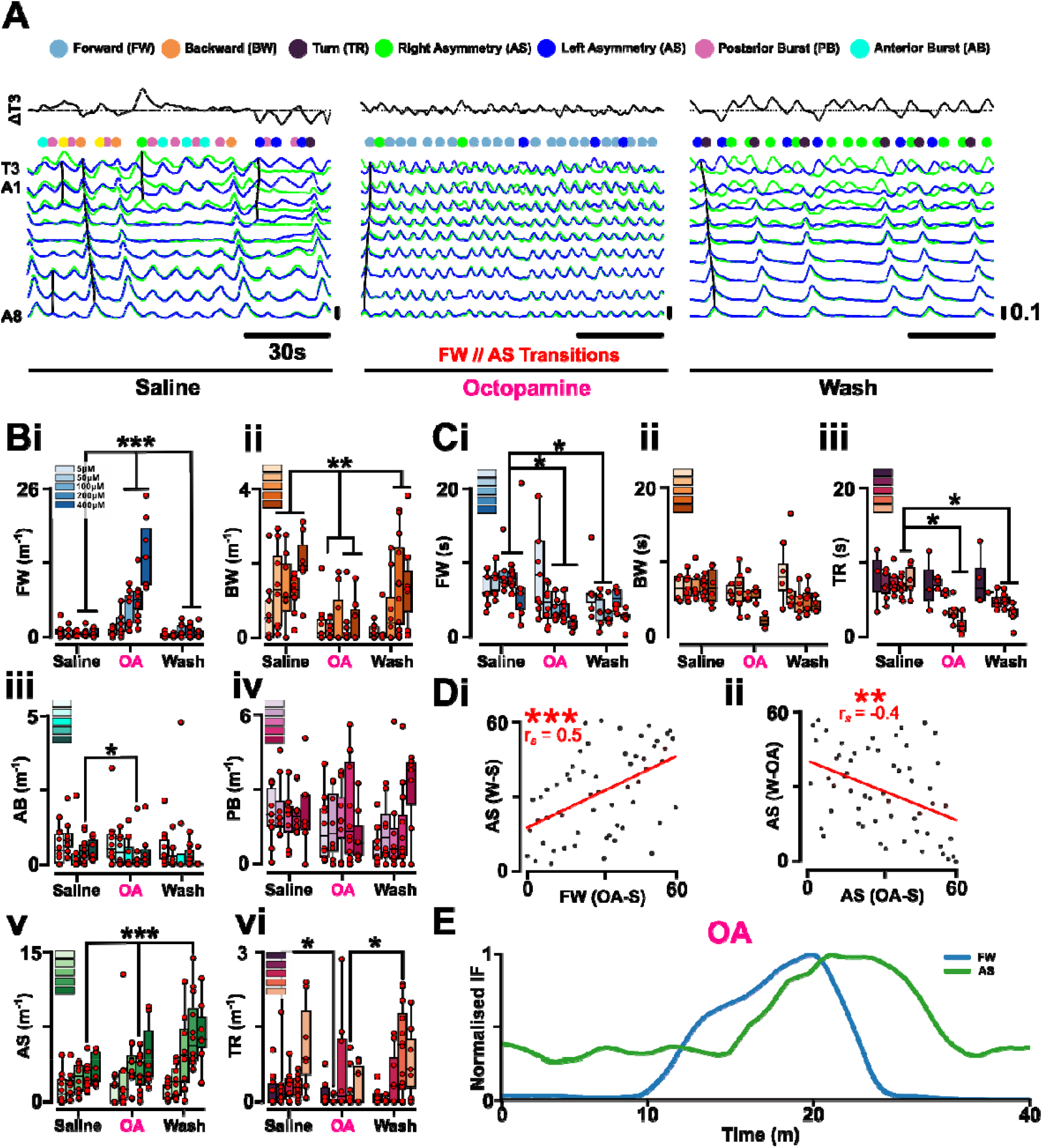
Bath Application of Octopamine Induces Fictive Forward Bias in the isolated larval CNS. **(A)** Exemplar calcium dynamics in left (blue) and right (green) motor neurons in the 3^rd^ instar isolated *Drosophila* ventral nerve cord (VNC) over 45-minute recording period under saline, octopamine (OA) (200μM), wash bath applications. Difference trace for T3 (left T3 – right T3 = ΔT3) shown above all traces. Dots above the traces indicate the initiation of the different fictive behaviours (light blue, fictive forward; orange, fictive backward; cyan, anterior symmetric burst; pink, posterior burst; blue, left anterior asymmetry; green, right anterior asymmetry). Black lines indicate the shape of each fictive activity to aid user’s recognition of fictive programmes. Note that OA application induces fictive forward bias with intermittent asymmetric activity. The post-application wash period exhibiting dominating anterior asymmetric activity. **(B)** Changes in fictive frequencies of different fictive behaviours under the application of OA concentrations (5 - 400μM) between saline, OA perfusion, and wash conditions for fictive: **(i)** forward, **(ii)** backward, **(iii)** anterior burst, **(iv)** posterior burst, **(v)** anterior asymmetric headsweeps, and **(vi)** turn. Note that the y-axis showing the rate of each fictive activity is different for each fictive activity. **(C)** Activity propagation speed from posterior (A8) to anterior (T3) for **(i)** fictive forward and anterior-to-posterior for **(ii)** fictive backward and **(iii)** fictive turning behaviours. **(Di)** Ranked relationship between the relative effect of OA on promoting fictive forward activity and the relative change in headsweep asymmetric. **(Dii)** Ranked relationship between the relative promotion of anterior asymmetric activity by OA and the change in anterior asymmetric activity in wash relative to OA application activity level. **(E)** Normalised instantaneous frequency (IF) of fictive forward (FW) and anterior asymmetric (AS) activity pooled across all 200μM OA preparations showing averaged elevated FW during OA application and average elevated AS in the post-application window. Post-hoc comparison were conducted with p-values: < .5 (*), < .01 (**), <.001 (***) shown. Only statistically significant bars are shown.

Exogenous bath application of OA increases intersegmental propagation for fictive forward and turn but not fictive backward activity (Figure 2C) similar to OA-inducing faster crawling in intact *Drosophila*. Fictive forward wave (Figure 2Ci) and turn wave (Figure 2Ciii) duration is significantly shorter during OA application. Specifically, all OA concentrations ≥50μM significantly increased forward wave speed (50 μM 4.4±2.4 s p = .05; 100 μM 3.6±1.3 s p < .001; 200 μM 3.7±1.4 s p = .002; 400 μM 1.7±0.7 s p = .01) compared to pre-application saline perfusion (8.4±8.3 s). Similarly, OA concentrations ≥200μM significantly increased turn wave speed (200 μM 3.2±1.6 s p = .01; 400 μM 1.6±1.4 s p = .03) compared to pre-application saline perfusion (7.0±1.9 s). OA did not induce any significant changes in fictive backward wave speed.

To understand the relationship between OA’s induction of fictive forward bias and the post-application anterior asymmetric fictive activity, we examined the relative changes in fictive frequency between different motor programs across all concentrations (Figure 2D). The more OA promoted fictive forward bias, the greater the post-application anterior asymmetric activity relative to pre-application (Figure 2Di, r_s_ = 0.5, p < .001). This proportional relationship was not exhibited between any other fictive events. The proportional increase in anterior asymmetric activity with respect to fictive forward bias appears significantly conditional on the relative promotion of anterior asymmetric activity during OA application (Figure 2Dii, r_s_ = -0.4, p = .002). The greater fictive forward bias promoted by OA, the more fictive anterior asymmetric activity is delayed until the network is no longer forward biased. During OA application, the induced fictive forward bias is regularly interspersed with a few anterior asymmetric events implying while OA potentiates the network towards fictive forward activity it does not completely suppress anterior asymmetric activity (Figure 2A, Octopamine, “FW // AS Transitions”). Aside from changes in the frequency of fictive events, OA induces a decrease in the amplitude of the calcium signal (saline: .22±.09; OA: .11±.05; wash: .13±.07 m^-1^, p < .001) without a significant change in signal baseline (Figure 2Aii, Figure 3A-E). The reduction in calcium amplitude was reflective of the reduced duration in motor bursting during OA application in both thoracic and abdominal motor pools (saline: 5.4±0.9s; OA: 3.8±1.8s; wash: 4.2±1.9s, p = .05) (Figure 3F-H). Taken all together, this implies OA induces fictive forward bias and potentiates anterior asymmetric activity, but the former dominates in isolated VNC output over the latter until OA’s removal removes the neuromodulatory potentiation of fictive forward activity.

**Figure 3.**
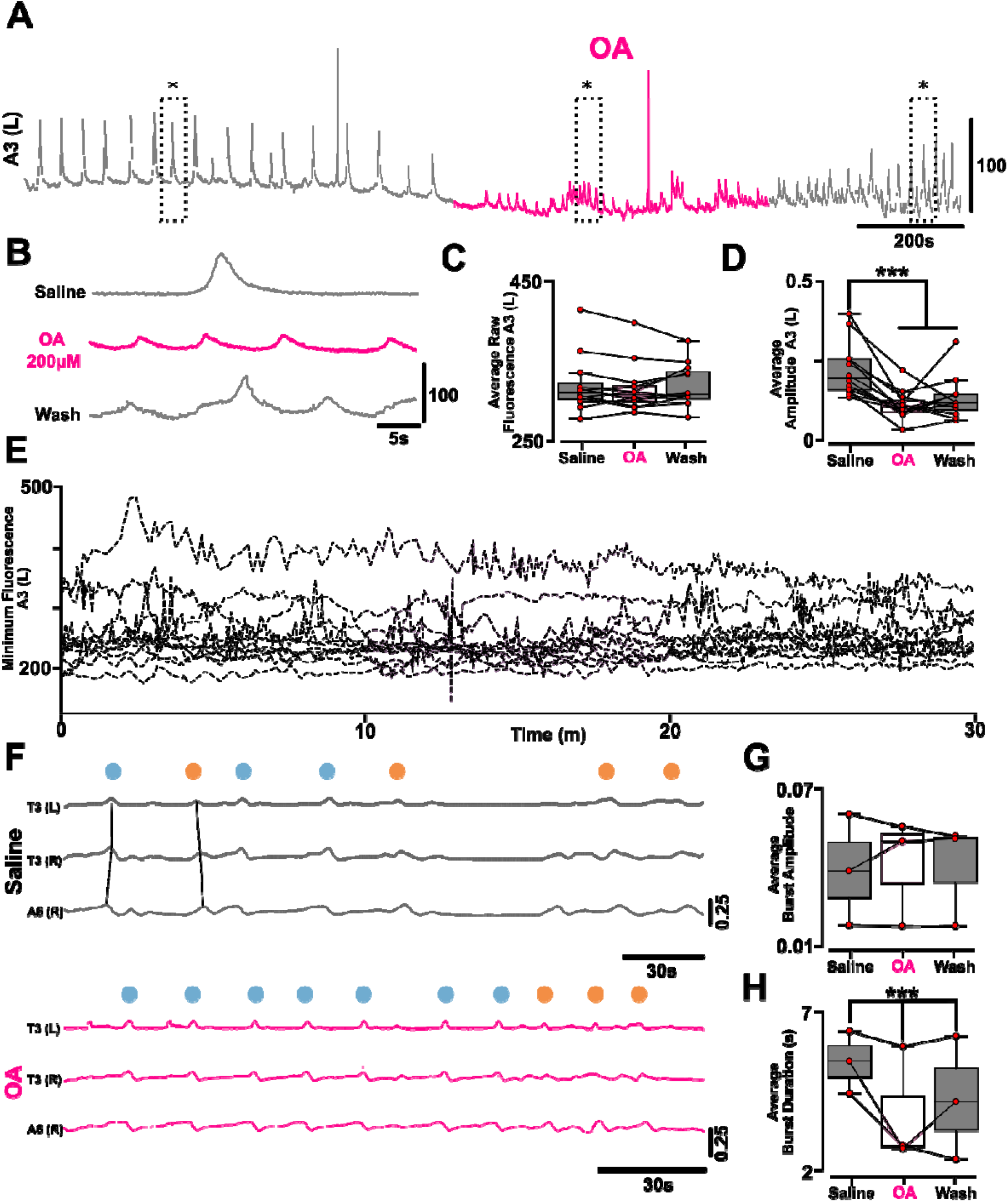
Octopamine Decreases Calcium Signal Amplitude and Motor Root Burst Duration Despite No Baseline Change. **(A)** Representative calcium signal trace from A3 left motor neuron before (blue), during 200μM octopamine application (magenta), and after (blue). **(B)** Exemplar activity in the left A3 calcium signal before, during 200μM octopamine application and after. Traces in B are subsections of the total trace denoted by *. **(C)** The average fluorescence value from calcium signal trace before, during, and after 200μM octopamine application. **(D)** The averaged peak amplitude from all A3 motor neuron calcium activity from calcium signal trace before, during, and after 200μM octopamine (OA) application across full ventral nerve cord (VNC) perfusion experiments (N=12). **(E)** All traces from full VNC perfusion experiments (grey, dashed line) before, during (magenta box), and after 200μM octopamine application. Note the absence of a visual change in baseline during octopamine application (magenta box). **(F)** Nerve root recordings for full VNC before (blue) and during (magenta) 200μM octopamine bath application. Yellow lines indicate fictive patterns. Note the elevated frequency of fictive forward wave activity during octopamine application (yellow lines, magenta traces). **(G)** The average amplitude of peaks in A6 motor neurons before and after 200μM octopamine application (N=3). **(H)** The average burst duration of peaks in A7 motor neurons before and after 200μM octopamine application (N=3). Note, the decreased burst duration in motor activity represents calcium activity as a lower signal amplitude. Post-hoc comparison were conducted with p-values: < .5 (*), < .01 (**), <.001 (***) shown. Only statistically significant bars are shown.

### TA Bath Application Induces Fictive Overlap & Motor Competition

We next evaluated the effect of TA on neuronal and fictive activity. Like the previous OA experiments, we exogenously bath applied TA onto isolated 3^rd^ instar *Drosophila* larva VNCs. TA did not induce a consistent fictive motor bias across applied concentrations compared to pre-application (Figure 4A, B). Across all fictive behavioural patterns, the only significant changes occurred post-TA wash. Specifically, fictive forward (200 μM 0.6±0.4 m^-1^ p < .001) and posterior burst (400 μM 0.7±0.6 m^-1^ p < .001) activity was significantly reduced in the wash period relative to the TA application period (FW, 200 μM 10.3±8.1 m^-1^) or pre-application saline period (PB, 200 μM 18.7±11.5 m^-1^) respectively. TA did not induce any significant changes to the intersegmental propagation of either fictive forward, backward, nor turn behaviours (Figure 4C), and did not affect the probability of transitions between any type of fictive event (Supplementary Figure 1C).

**Figure 4.**
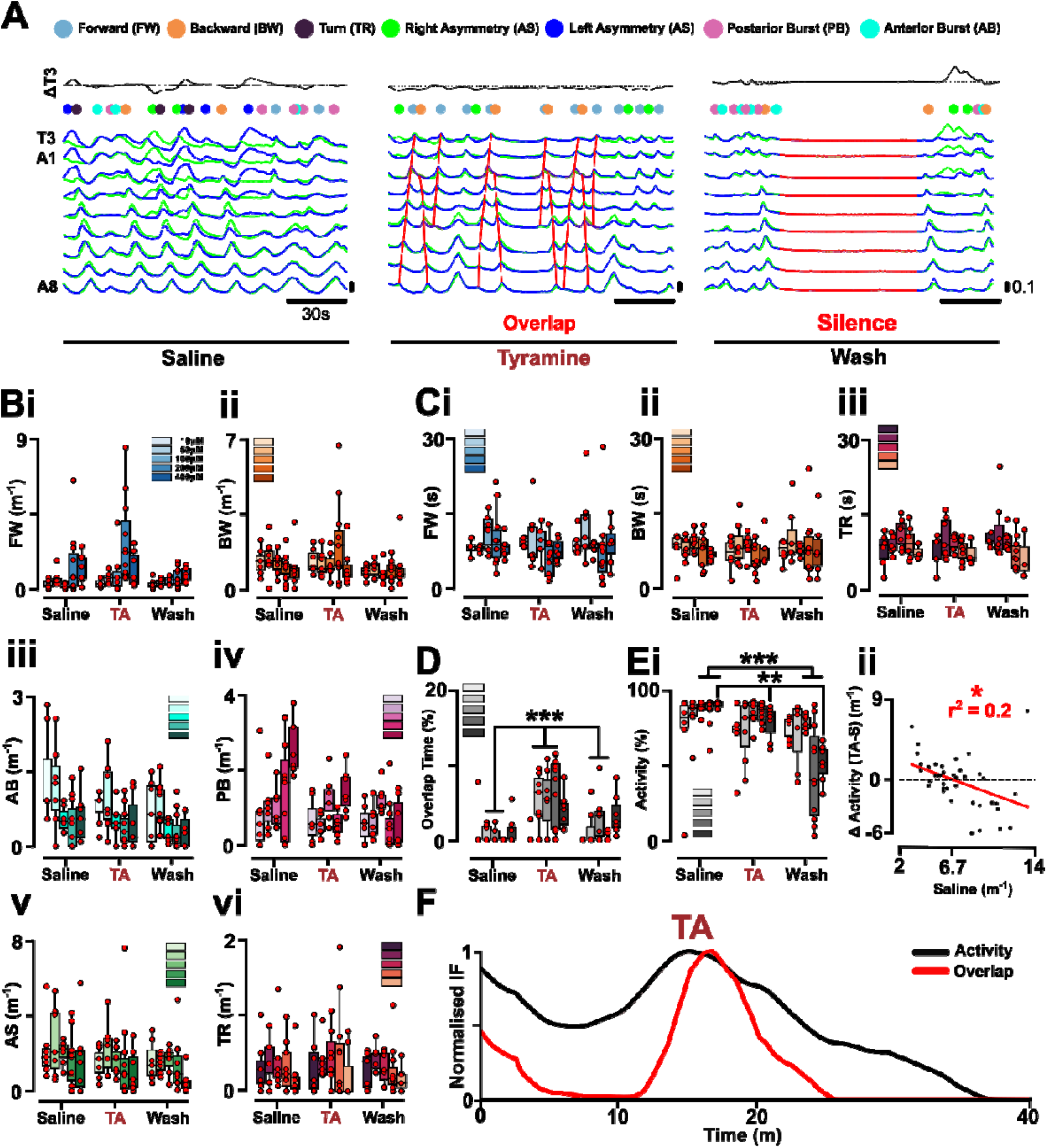
Bath Application of Tyramine Induces Fictive Motor Overlap and Competition. **(A)** Exemplar calcium dynamics in left (blue) and right (green) motor neurons in the 3rd instar isolated *Drosophila* ventral nerve cord (VNC) over 45 minutes under saline, tyramine (TA) (200μM), wash bath applications. Note the overlap between fictive forward and backward motor activity (red) during tyramine application and absence of fictive activity in the wash period. **(B)** Changes in fictive frequencies of different fictive behaviours under the application of TA concentrations (10 - 400μM) between saline, TA perfusion, and wash conditions for fictive: **(i)** forward, **(ii)** backward, **(iii)** anterior burst, **(iv)** posterior burst, **(v)** anterior asymmetric headsweeps, and **(vi)** turn. Note that the axis showing the rate of each fictive activity is different for each fictive activity. **(C)** Activity propagation speed from posterior (A8) to anterior (T3) for **(i)** fictive forward and anterior-to-posterior for **(ii)** fictive backward and **(iii)** fictive turning behaviours. **(D)** Percentage of time where overlap in activity between segments exists for pre-application, application, and post-application of TA across all concentration ranges. **(Ei)** Percentage of time where there is activity in at least one segment for pre-application, application, and post-application of TA across all concentration ranges. Note the concentration-dependent reduction in total activity within the wash period. **(Eii)** The relationship between overall pre-application fictive activity (“Saline Activity”) and the change in overall fictive activity during TA application (“TA vs Saline”). Note the activity-dependent inhibitory effect of tyramine. **(F)** Normalised instantaneous frequency (IF) of any and all fictive activity (black) and fictive program overlap (red) activity pooled across all 200μM TA preparations showing elevated activity and overlap during OA application and average suppression of activity in the post-application window. Post-hoc comparison were conducted with p-values: < .5 (*), < .01 (**), <.001 (***) shown. Only statistically significant bars are shown.

TA perfusion induced an increase in the overlap of fictive motor programs during application (Figure 4A, Tyramine “Overlap”) represented by wave activity commencing during the progression of a prior, incomplete fictive wave event. Perfusion of 200μM TA induced an increase in time the network demonstrates activity overlap across at least one segment (6.2±0.4 % p = < .001) compared to the 0% overlap demonstrated during the pre-tyramine application period (Figure 4D). The application of ≥100μM TA also induced a decrease in the activity in the post-application in a concentration-dependent manner (100 μM 84.5±4.8 % p = .01; 200 μM 49.8±31.7% p = .002; 400 μM 56.4±12.3 % p = .006) compared to pre-application period (100 μM 95.4±3.8%; 200μM 93.2±9.8%; 400 μM 97.7±2.2%) (Figure 4A, Wash “Silence”; Figure 4Ei). A consistent promotion of fictive overlapped events and post-application activity suppression is exhibited across pooled preparations (Figure 4F). Increasing the duration of the TA application window delayed the peak of any inhibitory silence effects, thus TA’s inhibitory effects are unlikely a product of delayed action but instead a product of network recovery post-TA’s neuromodulation (Supplementary Figure 2).

Given prior work has drawn attention to the state-dependent effects of TA inhibition, we next evaluated if TA’s effects on fictive activity were dependent upon the activity state. TA did exhibit proportional and increasing inhibitory effects as overall fictive activity levels rose beyond pre-application activity levels > 6.7m^-1^ (p = .04, r^2^ = 0.2; Figure 4Eii). Thus, when baseline fictive activity is low, TA can have excitatory effects and vice versa. Taken together, TA promotes overlaps between different fictive motor programs and demonstrates long-lasting activity-dependent inhibition of overall fictive activity post application.

Given that previous work has indicated an inhibitory role for TA at an individual cellular level [49], we evaluated the effect of TA perfusion on motor bursting activity. TA perfusion significantly increased bout motor root bursting (saline: 2.7±0.5m^-1^; TA: 7.4±0.6m^-1^, p = .05) (Figure 5Ai, ii). However, TA perfusion did not significantly affect bursting duration (Figure 5Aiii), overall duration of inactivity (Figure 5Aiv), bursting amplitude (Figure 5Av), nor tonic baseline amplitude (Figure 5Avi). 200μM TA application induced a reduction (p = .001) in the amplitude of calcium activity (Figure 4A, Figure 5Bi, iii) without any changes to the baseline calcium activity (Figure 5Bii, iv).

**Figure 5.**
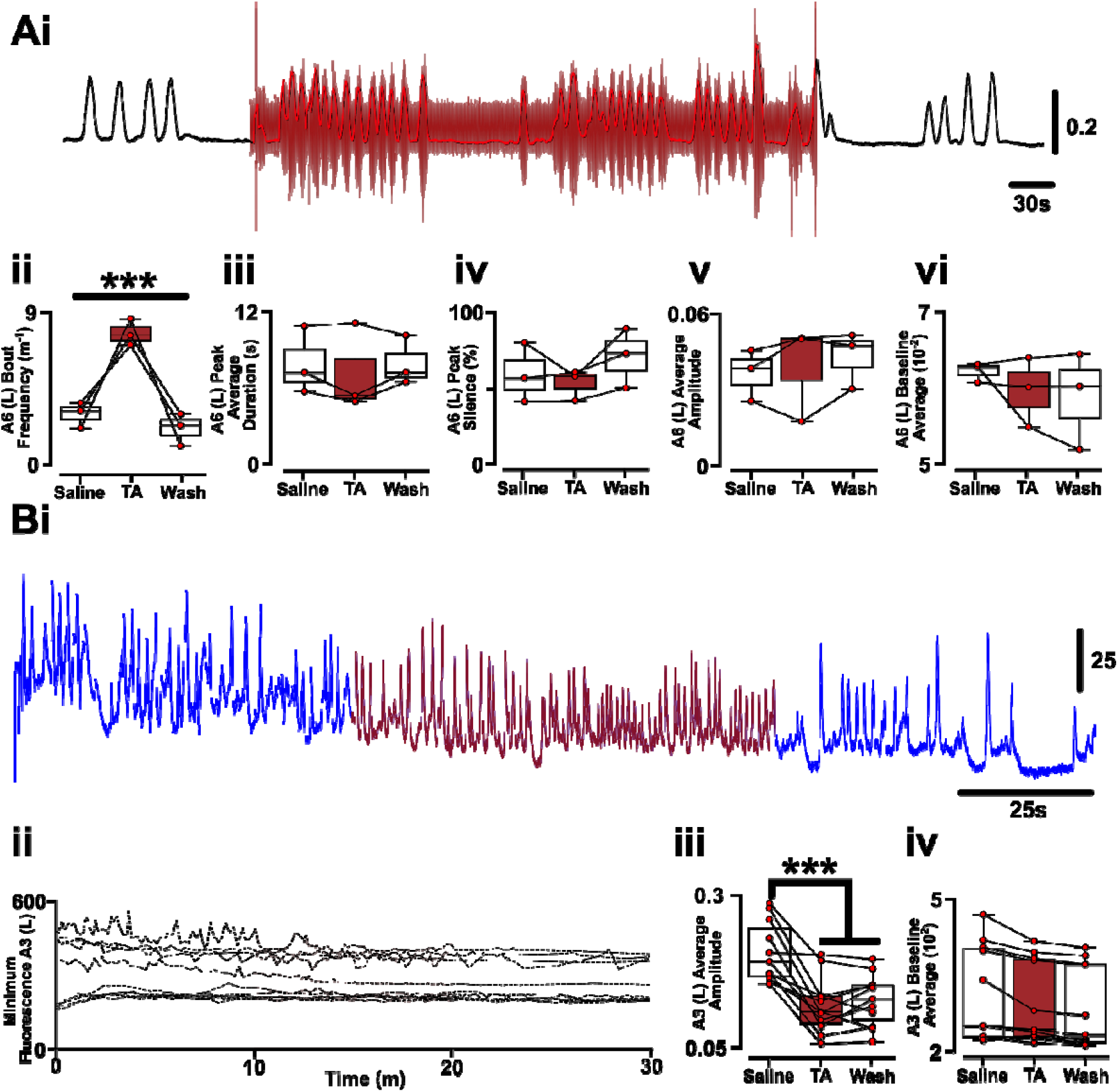
TA Shows No Inhibition of Motor Root Activity nor Calcium Baseline. **(A)** Effects of 50μM tyramine bath application on posterior abdominal motor nerve root activity demonstrating **(Ai)** subtle changes to firing patterns (N=3). The effect of 50μM tyramine bath application on posterior abdominal motor root **(Aii)** bursting frequency, **(Aiii)** average burst duration, **(Aiv)**, proportion of time (%) without activity > 5% change from baseline, **(Av)** average burst amplitude, and **(Avi)** baseline average for the pre-application (“Saline”), application (“TA”), and post-application (“Wash”) periods. (B) Representative effects of 200μM tyramine bath application (dark red) (N=11) on the (Bi) calcium activity on the motor neuron pool (abdominal segment 3, left: A3 (L)). **(Bii)** The baseline calcium fluorescence for all isolated ventral nerve cord preparations under 200μM tyramine bath application (dark red). The effect of 200μM tyramine bath application on **(Biii)** average calcium amplitude across fictive programs and (**Biv)** baseline calcium signal. Post-hoc comparison were conducted with p-values: < .5 (*), < .01 (**), <.001 (***) shown. Only statistically significant bars are shown.

### Simultaneous Co-application of Octopamine and Tyramine Promotes or Supresses Activity Based on Pre-Application Forward Bias

Given OA and TA conceptually act as antagonistic co-modulators in the intact system [45], we next evaluated the effects of simultaneous bath application of 200µM OA and TA on motor competition. Simultaneous bath application induced a bimodal effect on fictive activity level (p < .001). In one subset of preparations (N=8), simultaneous co-application retained fictive activity (saline: 11.2±8.6% vs OA/TA: 6.0±2.2% silence) whereas in another subset (N=11) co-application induced a collapse in distinguishable fictive rhythms (saline: 23.0±24.65% vs OA/TA: 82.17±13.27%, p = .004) which, like sole TA application, persists into the wash period (wash: 57.4±15.2%, p =. 007) (Figure 6Ai, Bi, C). However, within the preparations that exhibited a significant reduction in detected fictive rhythms, there persistent consistent dominant frequencies in calcium signals, consistent coherency between abdominal motor neuron pools, and consistent rhythmic organisation (see Method’s Analysis of Bimodality) (Supplementary Figure 3). Importantly, although combined OA and TA application elevated baseline fluorescence (saline: 176±21a.u. vs OA/TA: 185±20a.u., p = .01), peak event amplitudes under saline exceeded OA/TA baseline by 189±42a.u. (p <.001), indicating that dynamic range was preserved and OA/TA calcium signals were not compressed by tonic shifts in baseline (Supplementary Figure 4). Consequently, co-application of OA and TA either shifts the fictive network into a promotive mode or a low-amplitude, rhythmically-organised mode. Consequent analysis of fictive competition by co-application was performed on each cluster of the dataset independently.

**Figure 6.**
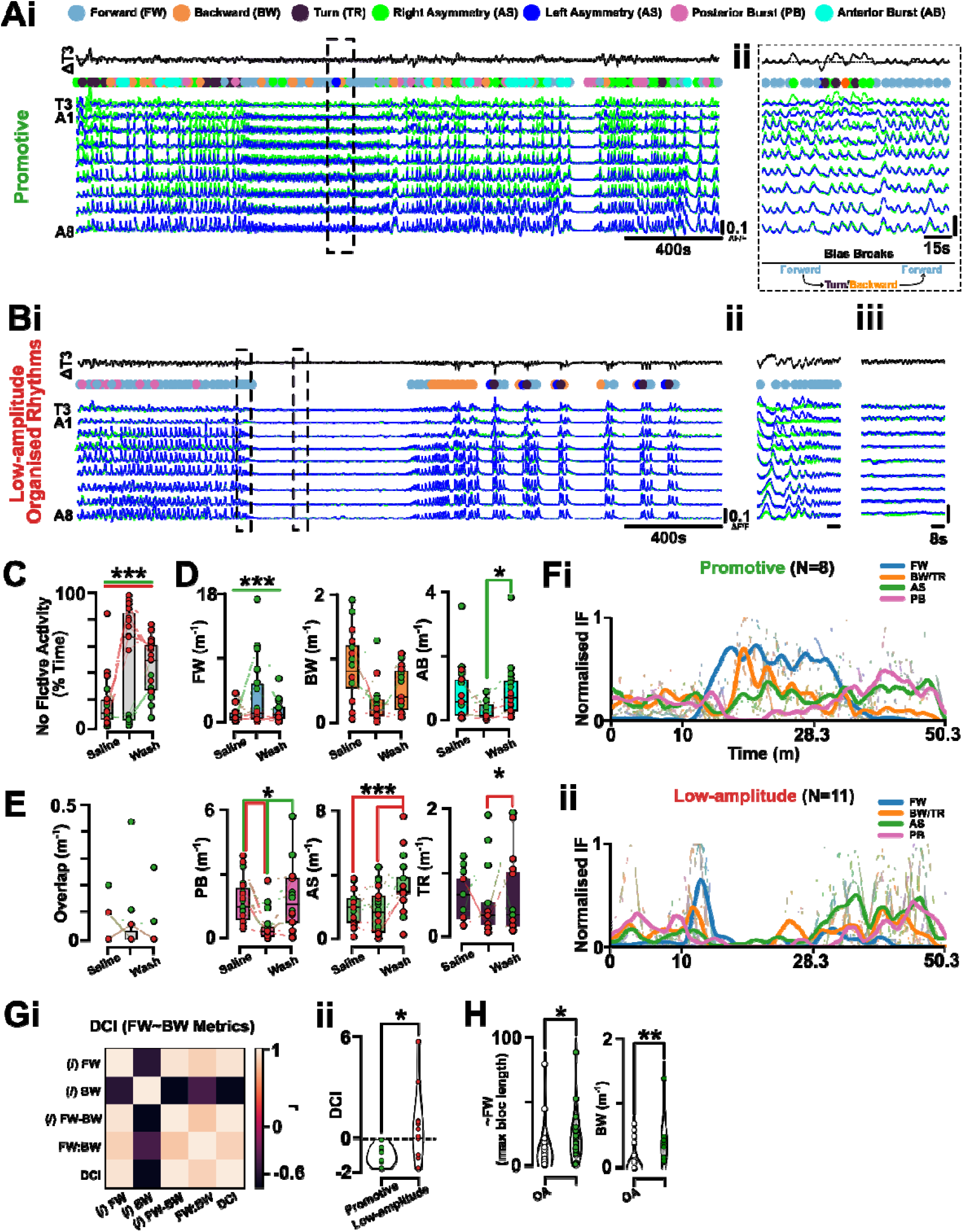
Simultaneous Perfusion Reveals Bimodal Effects on the Motor Competition Neuromodulatory Effects of OA and TA. **(Ai)** Representative calcium imaging trace for simultaneous perfusion of octopamine (OA) and tyramine (TA) that is either promotive (N=8) or induce low-amplitude organisation (N=11) of fictive activity. **(Aii)** Simultaneous perfusion induces fictive forward bias interspersed with breaks in bias consisting of fictive backward or turn activity. **(Bi)** Representative trace of OA/TA co-application inducing low-amplitude organisation of fictive rhythms with **(ii)** initial application inducing ellipsoid increases in fictive forward frequency which **(iii)** results in low-amplitude calcium signals with no distinguishable fictive rhythms. **(C)** The % of time isolated preparations display no determinable fictive activity (calcium activity peaks > 0.01 df/f) showing bimodality with preparations either showing no determined fictive activity (red) or promotion of activity (green) under simultaneous OA/TA application. **(C)** Changes in the frequency of fictive activity (forward = FW, backward = BW, anterior burst = AB, posterior burst = PB, anterior asymmetric = AS, turn = TR) between saline, simultaneous co-application of OA/TA, and wash period. **(D)** Changes in the frequency of overlapping fictive motor programs between saline, simultaneous co-application of OA/TA, and wash period. **(E)** Pooled and mean normalised instantaneous frequency (IF) of core fictive behaviours within the **(i)** promotive subcluster and **(ii)** suppressive subcluster upon simultaneous co-application of OA/TA. **(Fi)** Pearson’s correlation coefficient between the fractions of fictive forward and backward pre-application saline activity metrics showing correlation to the directional commitment index (DCI) with **(ii)** the one-dimensional forward vs backward fictive activity metric (DCI) sufficient to delineate promotive and low-amplitude, no determined fictive activity effects of OA/TA. **(H)** Changes of probability between fictive transitions between sole OA and combined OA/TA bath application. Post-hoc comparison were conducted with p-values: < .5 (*), < .01 (**), <.001 (***) shown. Only statistically significant bars are shown.

The promotion of fictive activity by simultaneous OA/TA bath application was not predominantly fictive forward and anterior asymmetric activity as exhibited in sole OA application, but instead blocs of fictive forward interspersed with fictive turn and backward motor programs (Figure 6Aii). However, compared against sole OA perfusion, OA/TA co-application did not increase the probability of bias-breaking blocs of activity, increase the number of blocs, nor increase the time spent in blocs. Instead, OA/TA co-application significantly increases the maximum length of consecutive non-forward activity (p = .01) within bias-breaking blocs (Figure 6H, left). In contrast, in preparations with overall suppression of detectable fictive activity consistently showed initial ellipsoid elevation in fictive forward frequency (Figure 6Bii) which evolved into low-amplitude calcium signals without distinguishable fictive rhythms (Figure 6Biii).

The co-application of OA and TA can differently affect the diversity of detectable fictive motor programs within and beyond the application period. During promotive neuromodulation, OA and TA co-modulation significantly increases the frequency of fictive forward activity (saline: 0.5±0.3m^-1^ vs OA/TA: 8.0±4.9m^-1^, p = 0.02) at the expense of posterior burst activity (saline: 1.9±0.6m^-1^ vs OA/TA: 0.3±0.3 m^-1^, p = 0.03) (Figure 6C). Interestingly, co-application of OA and TA resulted in a significantly higher frequency of fictive backward activity (OA: 0.1±0.2m^-1^, OA/TA: 0.5±0.4m^-1^, p - .001) than during OA application alone (Figure 6H, right). During neuromodulation that induced low amplitude organised rhythms, OA and TA co-modulation significantly decreased the frequency of detectable anterior asymmetric activity (saline: 1.7±1.1m^-1^ vs OA/TA: 1.1±1m^-1^, p = 0.01) and posterior burst activity (saline: 1.7±1.3m^-1^ vs OA/TA: 0.8m^-1^, p = 0.04). Interestingly, anterior asymmetric activity is uniquely promoted in the OA and TA co-modulation application period where low-amplitude organised rhythms are induced (wash: 3.2±2.0m^-1^, p = 0.01). Unlike with TA modulation, there was no significant change in the frequency of overlapping fictive motor programs during OA and TA co-application (Figure 6D). Overall, the simultaneous co-application of OA and TA either promotes fictive activity with both enhanced fictive forward activity – resembling sole OA application – alongside extended bouts of fictive backward activity without a post-application elevation in anterior asymmetric activity (Figure 6Fi) or collapses in detectable fictive activity into a low-amplitude, organised activity state which is accompanied by a post-application elevation in anterior asymmetric activity (Figure 6Fii).

By examining which features of the pre-application network were the strongest univariate discriminations, we found the effect of OA and TA co-application is conditional upon the pre-application forward-vs-backward state of the network. The pre-application switching rate between fictive motor programs, the level of activity silence, the diversity of transitions between states (i.e., entropy), nor the persistence of the network into any general fictive regime predicted evolution into promotive or low-amplitude rhythmic effects upon OA and TA co-application (Supplementary Table 2). Instead, directional dominance was exhibited solely by metrics associated with fictive forward and backward fictive behaviours. Univariate group comparisons (Mann–Whitney U tests with effect size estimation) and receiver operating characteristic (ROC) analyses demonstrated that forward– backward directional metrics exhibited the strongest discriminatory capacity between response regimes and are highly intercorrelated (r = 0.99 with FW−BW) (Figure 6Gi) (see Method’s Analysis of Bimodality). Given their high intercorrelation, we utilised principal component analysis to derive a single latent directional commitment axis (DCI). A threshold applied to the derived Directional Commitment Index correctly classified all promotive preparations and the majority of disorganised preparations (balanced accuracy ≈ 0.82) (Figure 6Gii). Specifically, isolated networks that exhibit forward-dominating activity exhibit collapse into a low-amplitude, organised motor regime under OA/TA co-application whereas backward-bias or liable networks exhibit activity promotion under OA/TA co-application (see representative Figure 6Ai, Bi dot annotations above traces).

### Sequential Perfusion of OA and TA Demonstrates Consistent Modulatory Effects

Given OA and TA demonstrate state-dependent effects, we next utilised each neuromodulator to bias the isolated network into different biased states and examined how the oppose neuromodulator shifts the fictive network in response. Here, we sequentially independently bath applied OA and then TA or vice versa and examined if there is a change in the underlying fictive dynamics of the isolated system.

Sequential application of OA and TA biased the isolated network akin to individual neuromodulator application. In either sequential configuration, application of 200µM OA biased the isolated network towards predominantly forward fictive behaviours (Figure 7A/Bi, OA) whereas 200µM TA induced an elevation in overlapping fictive events and long-lasting silence (Figure 7A/Bi, TA). Specifically, OA perfusion significantly elevated fictive forward frequency in both the OA-then-TA (saline: 1.0±0.4m^-1^, OA: 5.4±0.9m^-1^, p = .007, Figure 7Aii) and TA-then-OA (saline: 0.8±0.3m^-1^, OA: 4.5±1.1m^-1^, p = .04, Figure 7Bii) paradigm at the expensive of fictive backward behaviours. Application of TA induced significantly elevated increased frequency of overlapping fictive events uniquely after the OA perfusion window (saline: 0.03±0.03m^-1^, OA: 0.05±0.04m^-1^, TA: 0.69±0.14m^-1^, wash: 0.04±0.01m^-1^, saline vs TA: p = .05, TA vs wash: p = .05, Figure 7Aii) but, unexpectedly, not when initially perfused (Figure 7Biii). Similarly, the percentage of time the isolated network exhibited no detectable fictive activity was elevated and long-lasting by TA perfusion uniquely after the OA perfusion window (saline: 10.9±2.5%, OA: 7.8±3.4%, TA: 20.7±6.90%, wash: 38.4±6.6%, OA vs TA: p = .05, OA vs wash: p = .05, Figure 7Aiv), but not when initially perfused (Figure 7Biv). As exhibited by the normalized instantaneous frequency of fictive events, OA perfusion promotes more and faster fictive forward rhythms with trailing anterior asymmetric promotion whereas TA perfusion induces an increase in a diversity and general frequency of different motor programs (Figure 7Av, Bv). Thus, the order of sequential perfusion did not appear to alter the direct neuromodulatory effect of neither OA nor TA.

**Figure 7.**
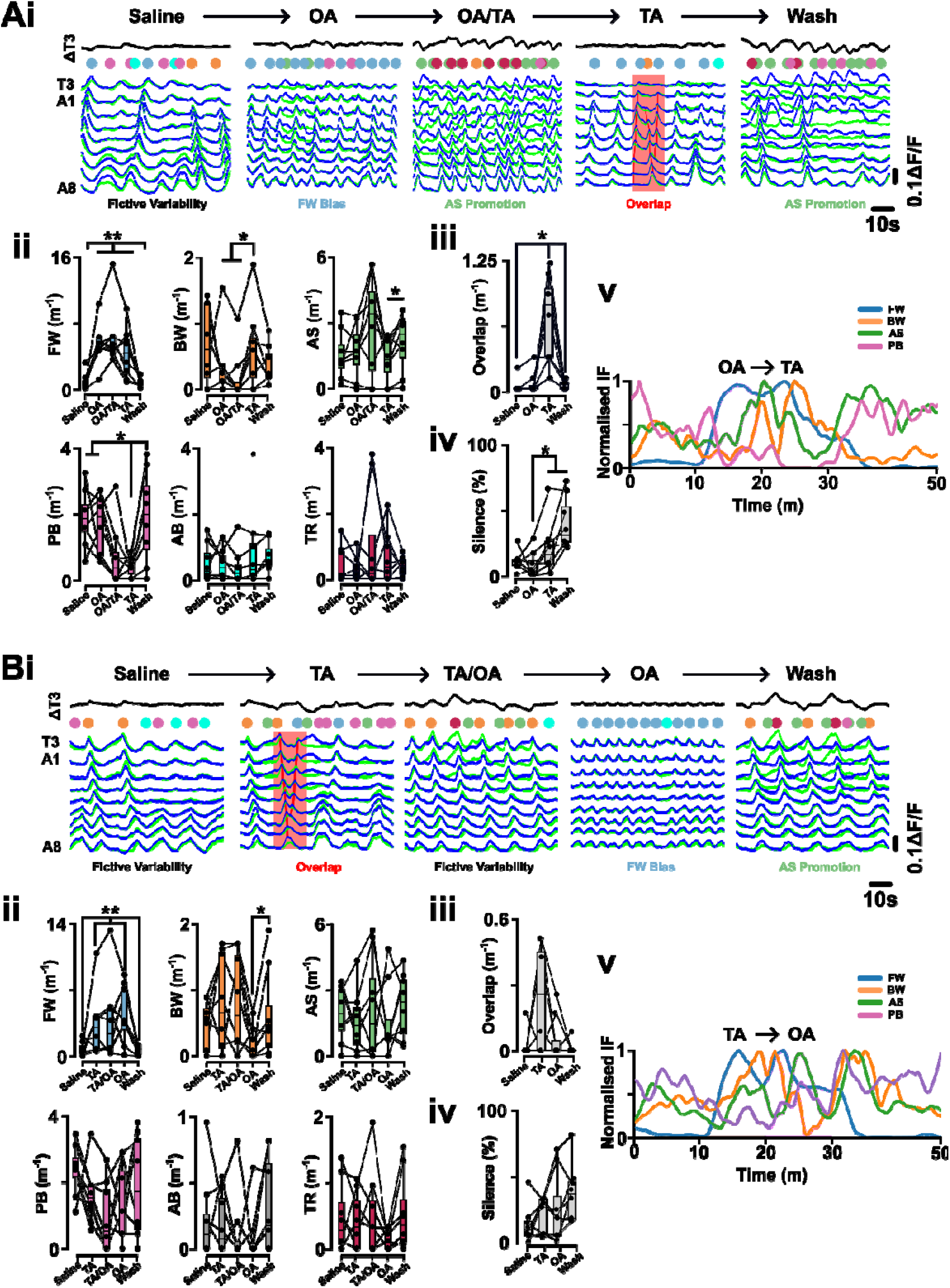
Sequential Perfusion Neither Changes the Motor Competition Neuromodulatory Effects of Octopamine nor Tyramine. **(A)** Sequential perfusion of octopamine (OA) followed by tyramine (TA) (N=8). **(Ai)** Representative calcium imaging samples from sequential perfusion of OA followed by TA. **(Aii)** Changes in fictive frequency (forward wave = FW, backward wave = BW, anterior asymmetry = AS, posterior burst = PB, anterior burst = AB, turn = TR). **(Aiii)** Changes in the frequency of overlapping motor programs (see “overlap” annotation in Ai). **(Aiv)** Change in the time (%) the fictive network exhibits no activity (i.e., Silence) (see annotations in Figure 4). **(v)** The evolving normalised instantaneous frequency (IF) of the significantly-varying fictive behaviours. (B) Sequential perfusion of tyramine (TA) followed by octopamine (OA) (N=7). **(Ai)** Representative calcium imaging samples from sequential perfusion of TA followed by OA. **(Aii)** Changes in fictive frequency (forward wave = FW, backward wave = BW, anterior asymmetry = AS, posterior burst = PB, anterior burst = AB, turn = TR). **(Aiii)** Changes in the frequency of overlapping motor programs (see “overlap” annotation in Ai). **(Aiv)** Change in the time (%) the fictive network exhibits no activity (i.e., Silence) (see annotations in Figure 4). **(v)** The evolving normalised instantaneous frequency (IF) of the significantly-varying fictive behaviours. Post-hoc comparison were conducted with p-values: < .5 (*), < .01 (**), <.001 (***) shown. Only statistically significant bars are shown.

## Discussion

Through acute drug manipulations, we demonstrate that the core adrenergic-like neuromodulators can dynamically alter the bias in a locomotory network’s output in a state-dependent manner with a by-product of extensive and sustained motor bias being an instantiation of alternate fictive activity after the end of a prior bias episode. Specifically, we demonstrated OA exhibits a bias-proportional promotion of an alternate mode – anterior asymmetric fictive activity – in response to forward-promoted bias. Furthermore, while TA is traditionally reported as an inhibitory neuromodulator, here we provided evidence for the activity-dependent promotive effects of tyramine on motor competition with increasing overlap events with inhibition dominating during wash periods. Co-application of TA and OA – more representative of intact adrenergic-like dynamics – revealed state-dependent neuromodulatory effects with co-application opposing the intrinsic bias within the pre-application network. A crucial element of sustained movement is action selection and the inhibition of those competing motor programs that induce different locomotion (e.g., forward vs backward). Here, we show how the isolated neural system appears to have an innate disposition to rebalance motor diversity by potentiating alternative motor patterns in response to suppression of motor diversity through sustained motor bias.

### Exogenous Application vs Intact Manipulations

The effects of exogenous application of octopamine (OA) and tyramine (TA) onto isolated preparations mirror the effects of genetic ablation and manipulations within intact animals. Specifically, elevation of TA levels either through TβH mutant *Drosophila* larva [21], [41] or feeding [49] induces dysrhythmic movement manifesting as slower movement with more pauses. Indeed, the inhibitory effects of tyramine are common to other invertebrates like *C. elegans* [28], [61]. Interestingly, here we showed exogenous TA application does not immediately induce pausing in activity but instead the induction of overlaps in fictive programs implying dysrhythmic, slower locomotion in intact *Drosophila* may not be a product of slower rhythm generation but incomplete motor generation that reduces movement speed and distance. In contrast, elevation of OA levels via direct application [15], [36], [74] or starvation [38], [41] are generally promotive of movement. Here, we found octopamine similarly induced elevated, fast fictive forward rhythms in line with behavioural studies. However, we found the sustained forward bias induced by octopamine also triggered long-lasting anterior asymmetric activity post application currently not exhibited in intact animal behaviour. In future, *Drosophila* behavioural studies could look at the long-lasting effects of sustained motor bias through acute manipulations as reported here.

### The Adrenergic-like System as a modulator of Motor Competition

Movement necessitates a balance between excitation-driven initiation of a rhythm and pattern generating circuit alongside inhibition of competing motor program-related networks. Motor competition - the dynamic interplay that underpins excitatory and inhibitory neural action - is one lens through which the OA/TA modulatory dichotomy has yet to be examined in detail.

The locomotor effects of modulating OA/TA signalling are well-reported across a variety of invertebrate species. Generally, OA/TA are considered mutually-antagonistic in locomotor neuromodulation with an excitatory/inhibitory binary [15], [44]. Complementing some studies that highlighted nuanced roles for adrenergic-like neuromodulation [44], we showed that TA exhibited activity-dependent inhibitory effects, and actually promoted overlapping between fictive programs, thus not leading to simplistic inhibitory effects. Indeed, decrements in locomotion exhibited when *Drosophila* larva have been fed TA [45] could well be failures to regulate motor competition rather than direct, general inhibitory effects. Those two competing explanations for TA effects are non-trivial differences. For instance, while TA has a hyperpolarising effect on an isolated neuron [49], we found an overall trend of TA to depolarise motor pool activity. Our indication that TA’s inhibitory effects may be a product of activity state can help marry the two bodies of thought. At a functional level, one therefore could consider OA/TA as mutual-antagonists altering motor competition rather than generally activity levels.

OA’s excitatory effect on fictive motor behaviour is reflective of an underlying inhibition of motor competition. Reminiscent of a large catalogue of prior studies [14], [21], [35], [43], [44], [45], we found exogenously-applied OA was excitatory by being promotive of forward fictive locomotion with higher intersegmental propagation of activity. Importantly, this promotion of locomotion, especially given the network appeared perpetually active with little to no episodes of non-activity, was reflective of significant inhibition of competing motor program occurrence. Indicative of a suppression of initiation, we here contextualise the neuromodulator OA, in comparison to TA, as a suppressor of motor competition. Reflective of a deeper role of the adrenergic-like system in motor competition, the emergence of proportional post-application activity changes due to OA’s neuromodulatory effects indicates a potential foundational connection in CPG competition. Given that asymmetric anterior activity can be considered a preliminary, and important element of backward locomotion [1], the fact OA’s induction of an exclusive fictive forward bias precipitated an elevation of such asymmetric anterior activity post-hoc suggests a potential underpinning dynamic in motor selection not previously appreciated. Our work points toward the possibility that a memory-like system may exist to compensate for extremes in fictive forward activity. Indeed, the emergence of post-application elevation in silence from TA perfusion may also be reflective of an opposing long-lasting effect following promotion of conflicting output. Unsurprisingly, inhibition is a vital element in regulating variability, as for a system to be able to devolve into different output modes, it must first have a configuration that allows it to sit at a potential junction of maximal variability (i.e., criticality) [1], [38], [75], [76], [77].

Co-application of OA and TA further highlights how the adrenergic-like modulation reveals important features of motor competition and intrinsic action selection. OA/TA’s co-application divergent effects suggest that intrinsic action of OA/TA is not inherently excitatory nor inhibitory, rather adrenergic-like neuromodulator act as gain-modulars whose effects depend on where the network sits along a forward-backward attractor axis before application. In the context of motor competition, it further implies that the evolution of a network’s output is non-random, but contingent upon prior bias state with networks intrinsically adaptive to respond to bias by promoting diversity. Overall, anterior asymmetric promotion in response to sustained bias, increased fictive inhibition in response to dysrhythmic overlapping activity patterns, and context-restrained promotion of fictive rhythms all highlight an unstudied, complex nature of neural systems.

### Conceptualising Neuromodulation of Motor Competition and Criticality

Ensuring appropriate movement is logically achievable by enabling motor networks to retain the ability to generate diverse range of motor programs while also transiently promoting one program over another. At a fundamental level, this dynamic interplay different CPG networks are competing in a winner-takes-all-like scenario is the essence of motor competition. The conceptualisation of network dynamics, competition between outputs, and the underlying dynamic interplay between excitation and inhibition as a source of output variability is well-documented in modelling and experimental studies of cortical network dynamics [78], [79], [80], [81]. An alternative way of interpreting our results is through the lenses of criticality [82], [83]. A ‘critical’ state is one in which a network can easily transition or ‘tip’ into other modes. For locomotory networks, being at a state of criticality would serve to ensure that the network is capable of easily transitioning to a the right motor program to suit the moment.

In this study, exogenous adrenergic-like manipulations reoriented the network away from producing a diversity of fictive motor programs into a restrictive fictive state: sustained neuromodulation pushed the CNS away from criticality towards a more biased state. For OA application, the engendering of a fictive forward bias state resulted in an anterior asymmetric state before return to baseline fictive diversity. For TA application, the induction of overlapping fictive behaviours – effectively increased unrestrained motor competition – induced bouts of silence before return to baseline fictive diversity. Thus, each neuromodulatory effect reveals potential different recovery pathways from sustained bias, back to a critical state. Indeed, through our acute manipulations, we found some indication of reorientation effects (e.g., anterior asymmetric promotion after OA-induced forward bias, prolonged fictive silence after TA-sustained fictive overlapping dysrhythmia) as the network reconfigures back to a state of being able to produce a diverse range of fictive patterns (i.e., the critical state). Other interpretations of the post-application effects of the neuromodulatory manipulations are possible. Plausibly, post-application effects may be a product of concentration differences: lower OA concentrations may preferentially potentiate anterior asymmetric activity rather than fictive forward activity. However, the potentiation of anterior asymmetric activity exclusively occurred post OA application, not as concentration was ramping up during application. Alternatively, differential expression of aminergic receptors subtypes may induce emergent effects on fictive rhythmogenesis [17], [21], [41], [42], [62], [84], [85] which could be interrogated in future genetic studies. Overall, our framework provides one possible explanation for how neural networks can produce a diversity of outputs that are both adaptable, and innately spontaneous. This work lays a foundation for exploring the underlying dynamics and memory of motor diversity within locomotor systems.

## Data Availability

The research data underpinning this publication can be accessed at https://doi.org/10.17630/9dd6da34-9064-4bff-b106-d32e4f4bf241.

## Author’s Contribution

S.R.P. and W.V.S. conceived the study. W.V.S conducted all experiments, analysis, and visualisation. W.V.S. and S.R.P finalised the figures. S.R.P. supervised the project.

## Acknowledgements

This project was made possible by an Industrial CASE PhD studentship (UKRI Biotechnology and Biological Sciences Research Council (BBSRC) grant number BB/T00875X/1) to WVS. This work was additionally supported in part by two Collaborative Research Grants awarded jointly by the Global Office of the University of St Andrews and The Halle Institute for Global Research at Emory University (awarded to S.R.P and A. Prinz), and The Global Office of the University of St Andrews and the University of Bonn (awarded to S.R.P and M. Pankratz).

## Supplementary Figures

**Supplementary Figure 1.**
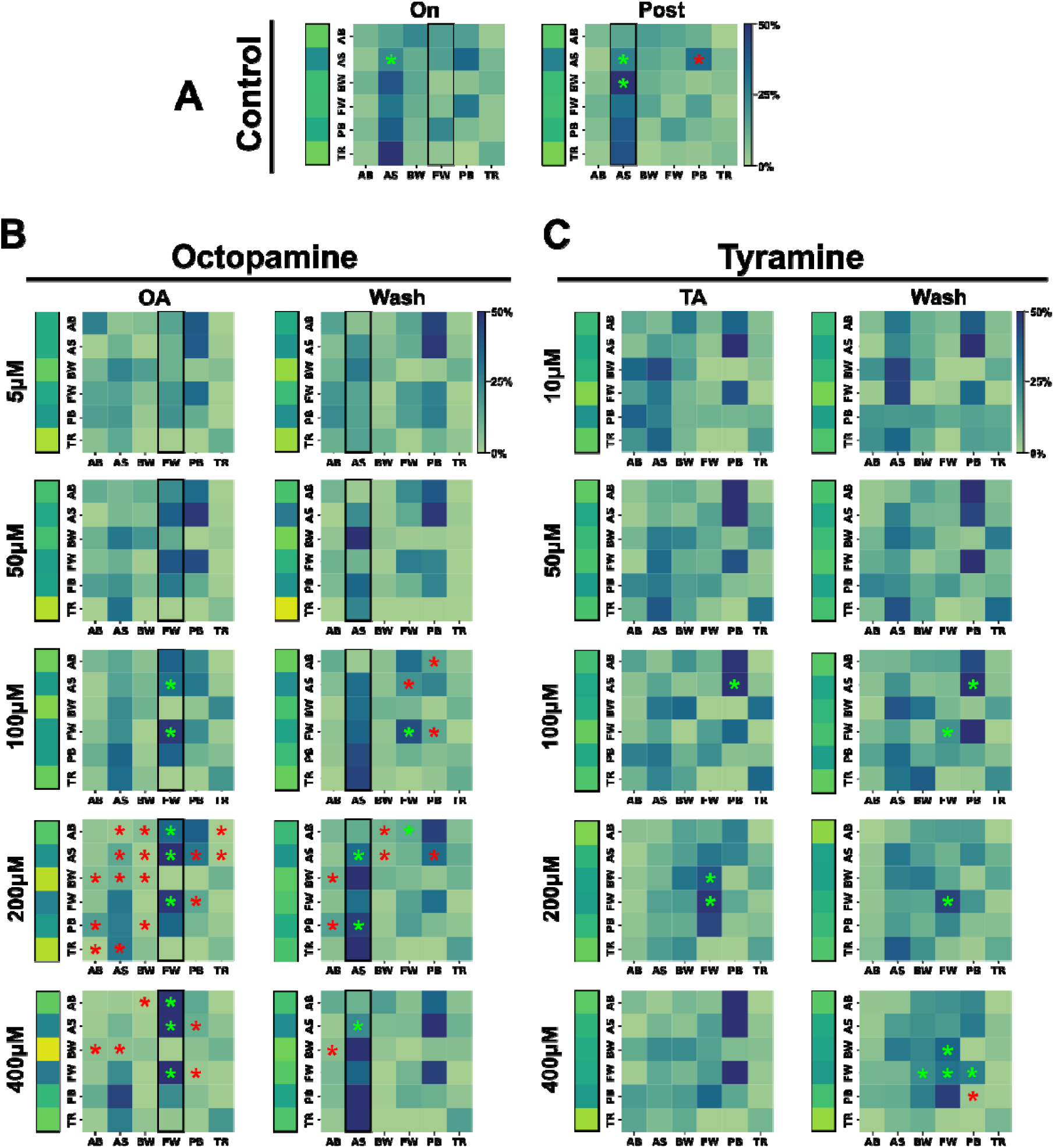
OA and TA Alters Motor Competition by Modulation Fictive Transition Probabilities. **(A)** The probability of any fictive behavioural transition (column) to a subsequent fictive program (row) within OK371>GCaMP6m control data (N=28) within the 10-to-20-minute time period (“On”) representative of the time period of the neuromodulator perfusion period and > 20-minute time period (“Post) representative of the time period of the neuromodulator wash period. “On” statistical comparisons made between the average transition probabilities in the 10– 20-minute period across all control isolated 3rd instar ventral nerve cords compared to the average transition probabilities in the 0–10-minute period. “Post” statistical comparisons are made between the average transition probabilities in the 20– 30-minute period across all control data compared to the average transition probabilities in the 0–10-minute period. **(B)** The average fictive transition probabilities across all the octopamine (OA) concentrations within the application (“On”) and post-application (“Wash”) period. **(C)** The average fictive transition probabilities across all the tyramine (TA) concentrations within the application (“On”) and post-application (“Wash”) period. Post-hoc comparison were conducted with p-values: < .5 (*), < .01 (**), <.001 (***) shown. Only statistically significant bars are shown.

**Supplementary Figure 2.**
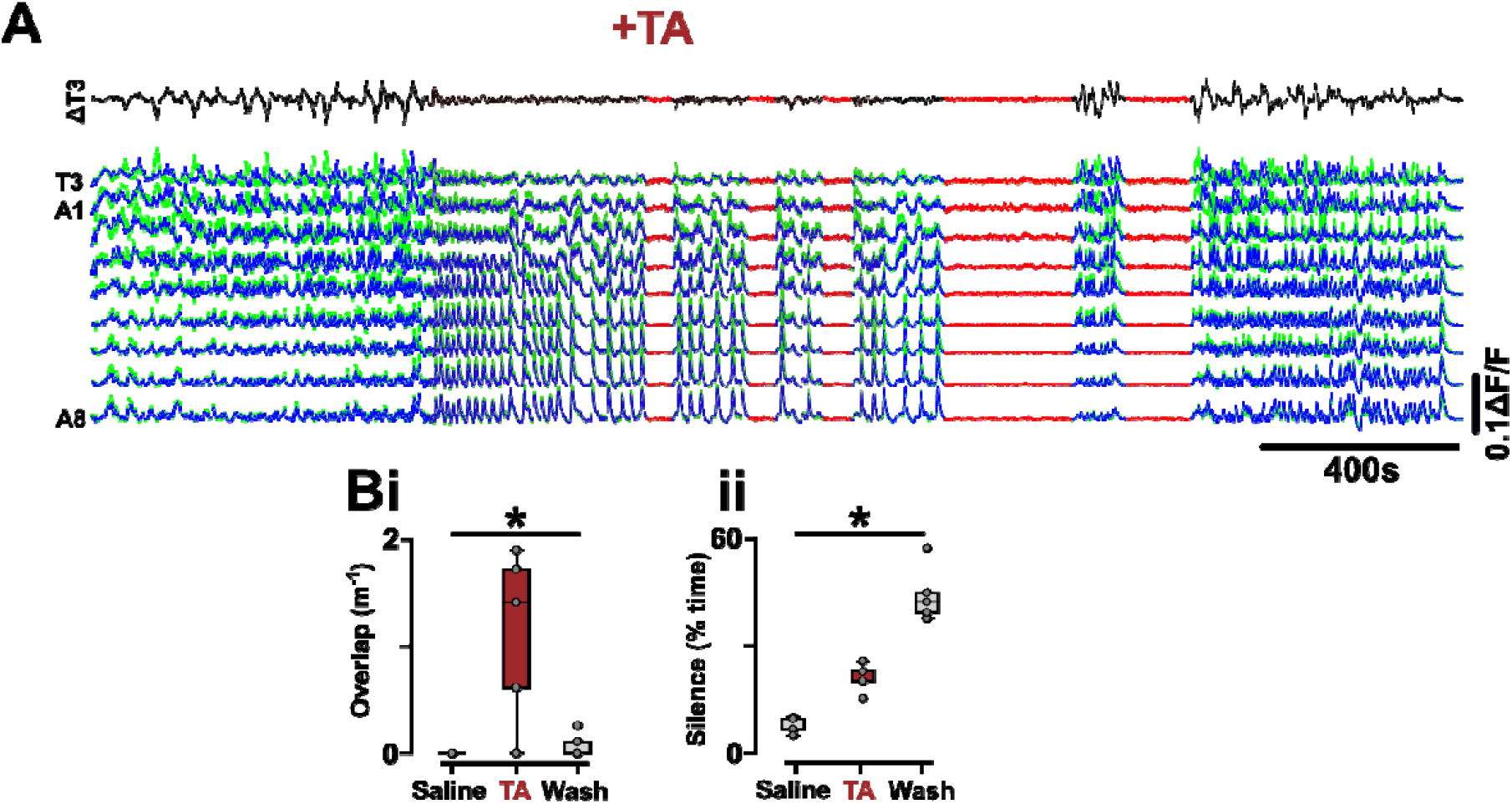
Long-lasting Tyraminergic Perfusion Demonstrates Post-Wash Inhibition is Unlikely a Feature of Slow-acting Inhibition. **(A)** Trace of long-lasting tyraminergic (TA) perfusion (18.3 minutes) showing silence (red) appears in the wash period despite the prolonged perfusion application window. **(B)** Long-lasting TA perfusion still induces **(i)** increased overlapping fictive motor programs and **(ii)** induced long-lasting inhibition of any glutamatergic motor neuron activity. Post-hoc comparison were conducted with p-values: < .5 (*), < .01 (**), <.001 (***) shown. Only statistically significant bars are shown.

**Supplementary Figure 3.**
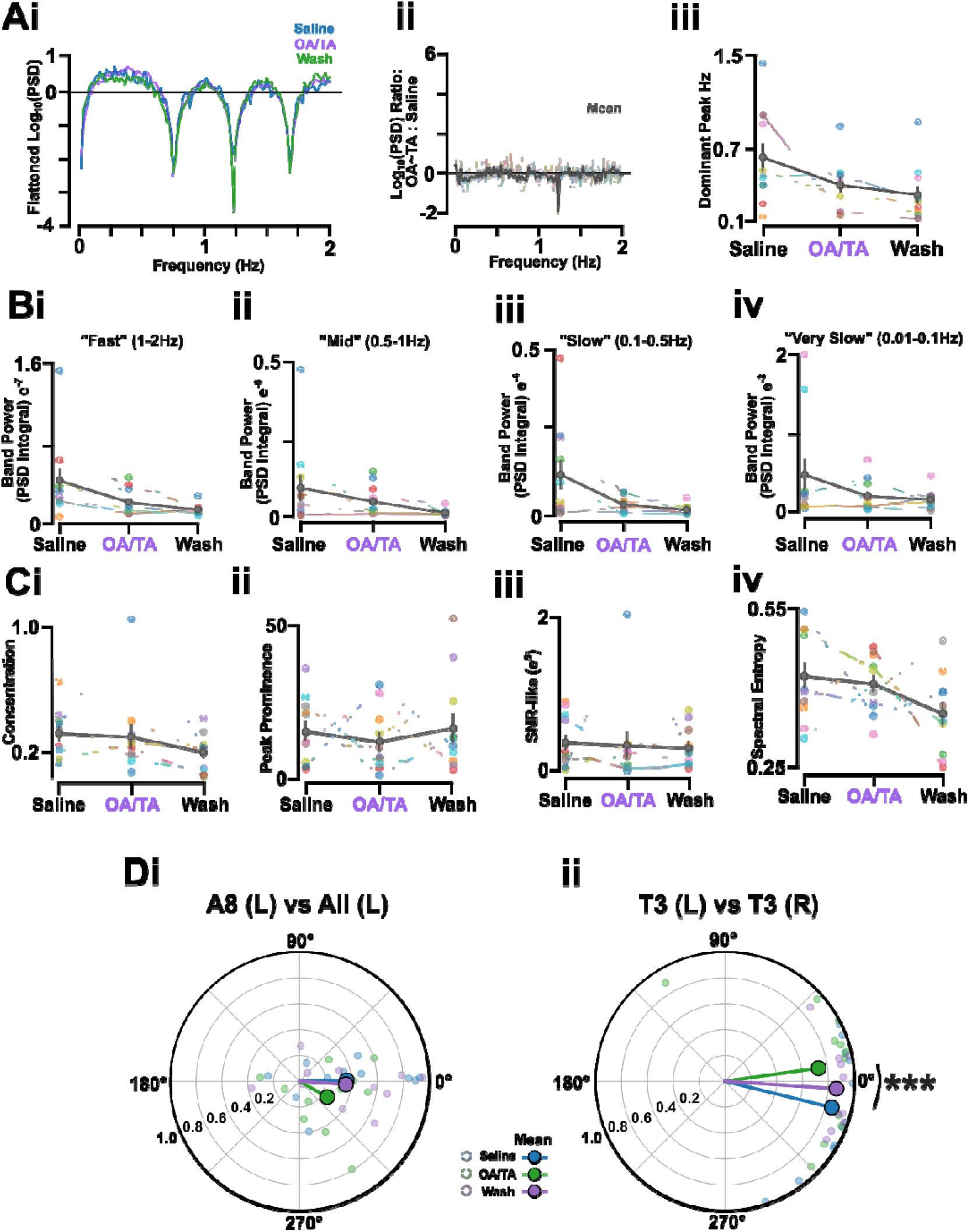
Power Spectra and Coherency of Low-Amplitude Calcium Signals During Octopamine & Tyramine Co-Application. **(A)** Power spectra density (PSD) for baseline-corrected averaged calcium trace for activity within saline, co-application of octopamine (OA) and tyramine (TA) and wash with **(i)** aperiodic flattening, **(ii)** ratio between OA/TA vs Saline, **(iii)** showing consistent dominant frequency. **(B)** The band power PSD for **(i)** fast (1-2Hz) frequency, **(ii)** medium (0.5-1Hz) frequency, **(iii)** slow (0.1-0.5Hz) frequency, and **(iv)** very slow (0.01-0.05Hz) frequency calcium signals in saline, OA/TA co-application, and wash demonstrating consistent oscillation frequency despite collapse in obvious fictive rhythms. **(C)** Metrics for quantifying structure in calcium signals – **(i)** concentration, **(ii)** peak prominence, **(iii)** signal-to-noise ratio-like (SNR-like), **(iv)** spectra entropy – demonstrate low-amplitude activity under OA/TA co-application unlikely induces disorganised activity in glutamatergic motor neuron pools. **(D)** Magnitude-squared coherence plots **(i)** between all left motor pools in abdominal (A1-7) and thoracic (T3) segments relative to the left A8 pool shows consistent phase relationship indicative of a similar profile of wave-like organisations albeit low-amplitude thus not detected by the Hill’s Valley Analysis algorithm and **(ii)** left vs right T3 motor neuron pools indicating a significant, consistent phase shift (10-15°) in left vs right T3 motor coordination, representative of persistence of anterior asymmetric activity in the OA/TA co-application period. Post-hoc comparison were conducted with p-values: < .5 (*), < .01 (**), <.001 (***) shown. Only statistically significant bars are shown.

**Supplementary Figure 4.**
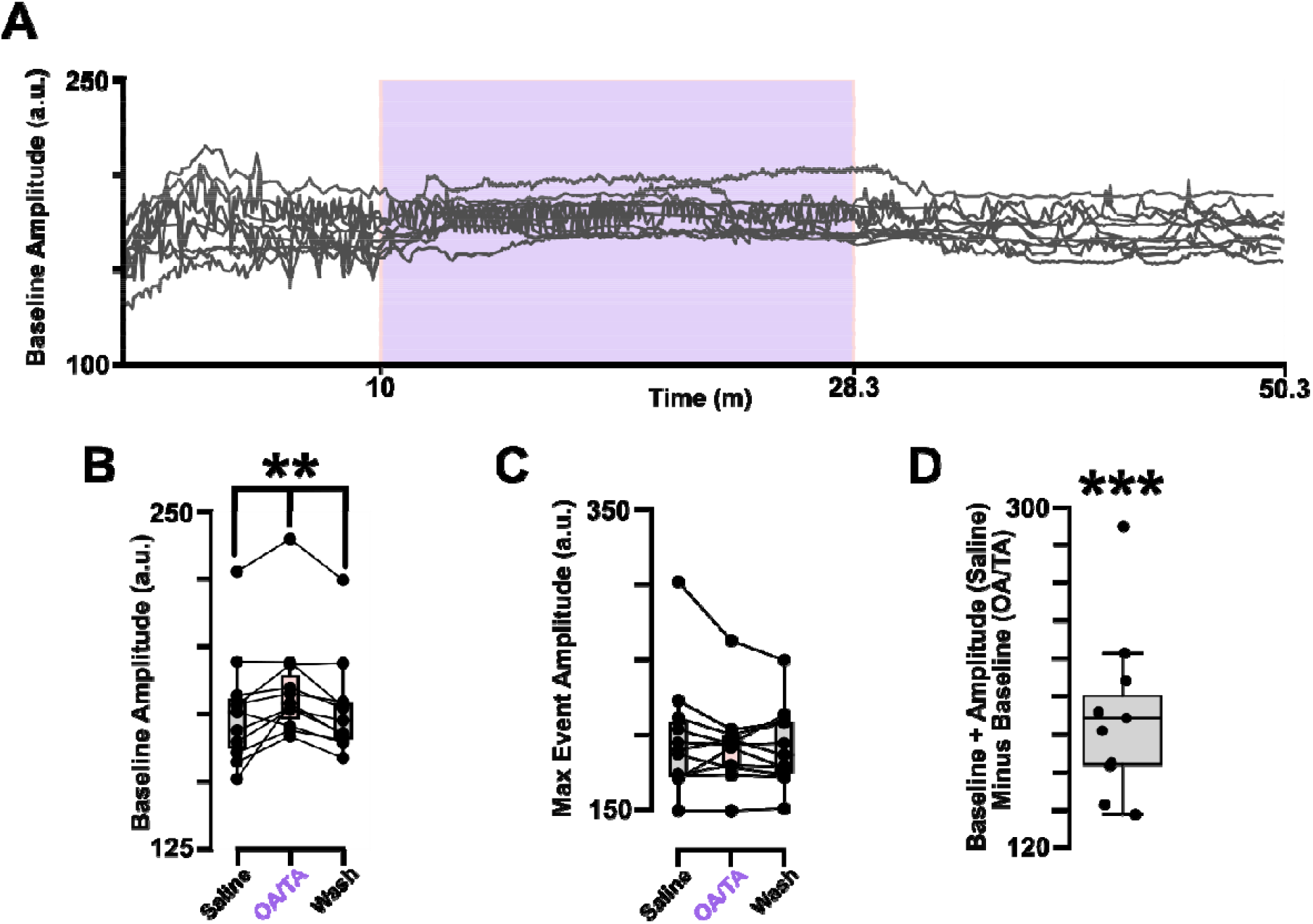
Calcium Indicator in Low-Amplitude OA/TA Co-modulation Preparations is Unlikely Saturated. **(A)** Raw A4 (L) calcium trace from isolated ventral nerve cord preparations (N=11) that exhibit suppressed fictive activity during OA/TA co-application. Red box indicates times when OA/TA is co-applied to the preparation. **(B)** The baseline amplitude of A4 (L) raw calcium signal. **(C)** The maximum amplitude of peaks in calcium activity in isolated nerve cords that exhibit suppressed activity upon OA/TA co-application. **(D)** The relative difference between the maximum peak amplitude in saline for suppressed prep and the changing in tonic calcium signal baseline indicating, despite elevated tonic baseline under OA/TA co-application, the dynamic range of the calcium signal still remains. Post-hoc comparison were conducted with p-values: < .5 (*), < .01 (**), <.001 (***) shown. Only statistically significant bars are shown.

**Supplementary Table 2.** Predictive Significance of OA/TA Pre-Application Fictive Features.

| Feature | p-value | Cliff's Delta | AUC |
| --- | --- | --- | --- |
| FW/BW | .012 | -.68 | .84 |
| FW%-BW% | .020 | .02 | .82 |
| BW% | .023 | .64 | .18 |
| AS% | .840 | .07 | .47 |
| FW% | .033 | -.59 | .80 |
| AB% | .041 | .57 | .22 |
| TR% | .049 | .55 | .23 |
| Saline Silence % | .310 | -.30 | .65 |
| PB% | .657 | -.14 | .57 |
| Switch Rate | .090 | .48 | .26 |
| Stickiness | .206 | -.36 | .68 |
| Transition Entropy | .033 | .59 | .20 |

